# Identification and Targeting of Cortical Ensembles

**DOI:** 10.1101/226514

**Authors:** Luis Carrillo-Reid, Shuting Han, Ekaterina Taralova, Tony Jebara, Rafael Yuste

## Abstract

Breaking the neural code requires the characterization of physiological and behavioral correlates of neuronal ensemble activity. To understand how the emergent properties of neuronal ensembles allow an internal representation of the external world, it is necessary to generate empirically grounded models that fully capture ensemble dynamics. We used machine learning techniques, often applied in big data pattern recognition, to identify and target cortical ensembles from mouse primary visual cortex *in vivo* leveraging recent developments in optical techniques that allowed the simultaneous recording and manipulation of neuronal ensembles with single-cell precision. Conditional random fields (CRFs) allowed us not only to identify cortical ensembles representing visual stimuli, but also to individually target neurons that are functionally key for pattern completion. These results represent the proof-of-principle that machine learning techniques could be used to design close-loop behavioral experiments that involve the precise manipulation of functional cortical ensembles.

## Introduction

The coordinated firing of neuronal populations is considered to be the substrate of sensory, behavioral and cognitive functions. Coactive neuronal groups, defined as neuronal ensembles, are assumed to generate complex circuit functions (Buzsaki, 2010; Cossart et al., 2003; Hebb, 1949; Luczak et al., 2009; Luczak et al., 2007; Mao et al., 2001; Miller et al., 2014). Recent advances in two-photon calcium imaging and two-photon optogenetics have made possible the simultaneous reading and writing of cortical ensemble activity with single cell resolution in awake animals (Carrillo-Reid et al., 2016). However, how the activation of specific groups of neurons relates to functional or behavioral changes has been difficult to elucidate, because it requires online identification of individual neurons that can be targeted during optogenetic experiments.

Cortical ensembles in primary visual cortex consist of coactive neurons (Carrillo-Reid et al., 2016; Cossart et al., 2003; Ko et al., 2011; Mao et al., 2001; Miller et al., 2014), forming a network structure that can be intuitively characterized with probabilistic graphical models, where nodes and edges are biologically meaningful, representing neurons and their functional connections respectively. Here, we exploit probabilistic graphical models to analyze two-photon calcium imaging recordings of mouse primary visual cortex. More specifically, we consider CRFs (Koller and Friedman, 2009), a combination of graph theory and probabilistic modeling. CRFs use graphs to simplify and express the conditional independence structure between a collection of random variables. Graph theory techniques have been used in neuroscience to describe the structural and functional organization of entire brains (Bullmore and Sporns, 2009). However, such graphs are usually constructed with nodes representing brain regions (He et al., 2007), and edges representing information flow (Iturria-Medina et al., 2008). For functional analysis, many studies have constructed graphs with data from fMRI, EEG and electrode arrays, taking brain regions (Achard and Bullmore, 2007; Fair et al., 2008; Hagmann et al., 2008), voxels (Eguiluz et al., 2005; van den Heuvel et al., 2008; Zuo et al., 2012) or electrode position (Downes et al., 2012) as nodes, and activity associations such as cross correlation, mutual information and Granger causality as edges (Bullmore and Sporns, 2009; Fair et al., 2008; Khazaee et al., 2015; Micheloyannis et al., 2009; Wang et al., 2010). In addition, at the single cell level, graphs have been used to describe organizing principles of artificial neural networks (Iturria-Medina et al., 2008; Sporns, 2000). Such graphs are usually associated with a restricted set of parameters that describe the weight and direction of edges obtained by pairwise metrics, therefore have limitations for characterizing changes in overall network dynamics and properties underlying population activity. Finally, a few studies have applied graph theory to describe network organization in calcium imaging data with single cell resolution in cultures or brain slices (Bonifazi et al., 2009; Gururangan et al., 2014; Yatsenko et al., 2015). Nevertheless, these methods have been applied only as a descriptive tool that lacks for a model that can be used to identify and manipulate with high spatial precision neurons that could have a potential role orchestrating the overall network activity in awake animals.

We demonstrate that CRF models allow not only the identification of cortical ensembles associated with different experimental and physiological conditions but also the prediction of the neurons that are most efficient at pattern completion. This method opens the possibility of targeting, with single cell precision, neurons from specific populations, to specifically alter microcircuit function. Importantly, different from previously used methods for neuronal ensemble identification, CRFs generate a functional model of the circuit that can be used to design and explore close-loop behavioral experiments.

## Results

### CRF models can encode population responses of visual cortex to oriented stimuli

CRFs model the conditional distribution *p*(**y**|**x**) of a network, where **x** represents observations and **y** represents true labels associated with a graphical structure (Sutton and McCallum, 2012). Since no assumptions are made on **x**, CRFs can accurately describe the conditional distribution with complex dependencies in observation variables associated with a graphical structure that is used to constrain the interdependencies between labels. Therefore, CRFs have been successfully applied in diverse areas of machine learning such as analysis of texts (Peng et al., 2011), bioinformatics (Li et al., 2008; Liu et al., 2006; Sato and Sakakibara, 2005), computer vision (He et al., 2004; Sminchisescu et al., 2006) and natural language processing (Choi et al., 2005; Lafferty et al., 2001).

In order to study the network dynamics of cortical ensembles we constructed CRF models using population responses to visual stimuli from layer 2/3 neurons of primary visual cortex in awake head-fixed mice (Figure 1A). Population vectors representing the coordinated activity of neuronal groups were inferred from calcium imaging recordings (Carrillo-Reid et al., 2015b) and used as training data (Figure 1B). We defined activity events from each neuron as nodes in an undirected graph, where each node can have two values: ‘0’ corresponding to non-activity, and ‘1’ corresponding to neuronal activity. In this way nodes representing neurons interact with each other through connecting edges, which have four possible configurations ‘00’, ‘01’, ‘10’, and ‘11’, depending on the values of two adjacent nodes. The two values associated with nodes and the four values associated with edges are characterized by a set of parameters called node potentials (*φ*_0_, *φ*_1_) and edge potentials (*φ*_00_,*φ*_01_,*φ*_i0_,*φ*_11_) respectively (Figure 1C). These parameters are also known as potential functions and reflect the scores of individual values on each node and edge. Using part of the observation data (e.g. training data), we estimated the model’s structure (e.g. graph connectivity) and parameters (e.g. the potentials) and then performed cross-validation on held-out data (Experimental Procedures). Once the model is learned from the data, the normalized product of the corresponding nodes and edge potentials describes the probability distribution of a neuronal population that represents a specific activation pattern.

**Figure 1.**
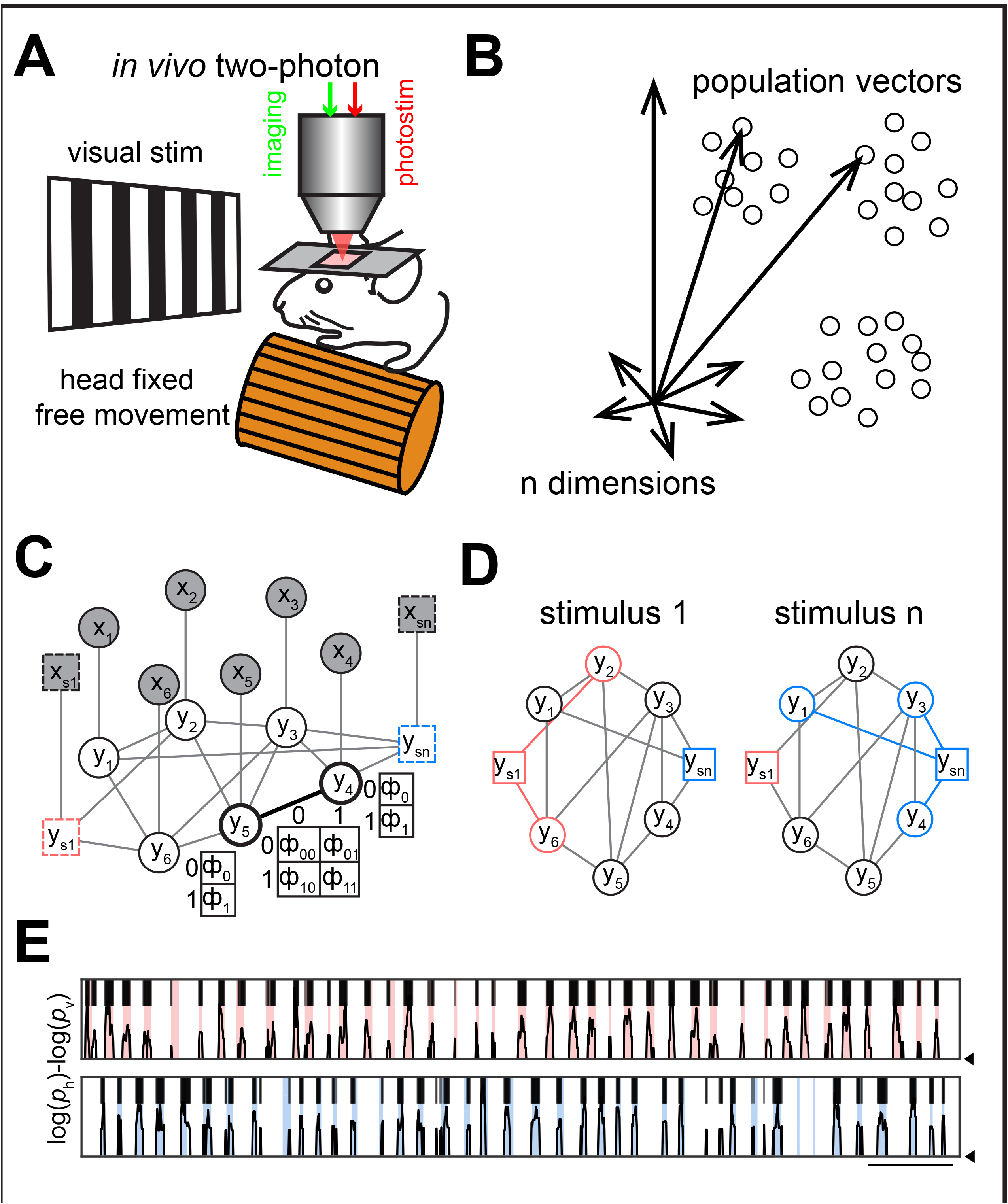
Classification of visual stimuli from calcium imaging data using CRFs. **A.** Experimental setup: simultaneous two-photon imaging and two-photon optogenetics were performed in layer 2/3 of primary visual cortex in head fixed freely moving mice. **B.** Schematic representation of population vectors. Each point in a multidimensional space represents a population vector defined by a coactive group of neurons. A neuronal ensemble is defined by a cluster of population vectors. **C.** Graphical representation of CRFs. Circles represent neurons. Squares represent added nodes depicting visual stimuli. Shaded nodes (x) represent observed data. White nodes (y) represent true states of the neurons, and are connected by edges that indicate their mutual dependencies. Node potentials are defined over the two possible states of each node, and edge potentials are defined over the four possible states of each existing edge, depending on the state of adjacent nodes. **D.** Graphical representation of isomorphic graphs highlighting cortical ensembles extracted from CRF models. Colors represent ensembles related to different stimuli. **E.** Ratio of log likelihood predicting horizontal (red; top) and vertical (blue; bottom) drifting-gratings calculated by CRFs. Colored stripes indicate visual stimuli. Scale bar: 10 seconds.

To integrate information about the external stimulus along with the observed neuronal data, we added an additional node for each type of stimulus that was presented to the animal. This node was set to ‘1’ when the corresponding stimulus was on and ‘0’ when the stimulus was off (Figure 1C). The general and mathematical properties of CRF models obtained with added nodes did not significantly differ from CRF models obtained without added nodes (Figure S1). In both conditions, CRFs modeled the conditional probability of network states given the observations. Therefore, by treating visual stimuli as added nodes and comparing the output likelihood of observing each stimulus, CRFs were able to model visual stimuli from observed data. In this way, the neurons directly connected to the added nodes represent a specific model for different visual stimuli (Figure 1D). Given two different visual stimuli (e.g. horizontal or vertical drifting gratings) and assuming that the probability (*a priori*) of visual stimuli is uniform, the ratio of the probability to observe each stimulus, taken from the model, can be used to classify presented stimuli (Figure 1E). These results demonstrated the ability of CRFs to accurately model different orientations of drifting-gratings from calcium imaging population recordings.

### Identification of core ensembles from CRF models

To identify core ensembles defined as groups of neurons that can efficiently represent different physiological functions (Carrillo-Reid et al., 2015b; Sadovsky and MacLean, 2014) we used the whole model obtained from CRFs and pruned the neurons that didn’t contribute to the efficient identification of visual stimuli. In order to do that we set the activity of each neuron to be either ‘1’ or ‘0’ in all population activity vectors of the dataset, and compared the probability ratio (logarithm probability difference) from the inferred CRF models (Figure 2A). Then, we calculated single neuron performance by binarizing the logarithm probability difference (Figure 2B) and calculating the area under the curve (AUC) from the receiver operating characteristic curve (ROC). Since cortical core ensembles have concomitant activity, we computed the node strength as the summation of edge potentials from active adjacent nodes (*φ*_11_ terms). In this way, high node strengths revealed recurrent coactive groups of neurons whereas low node strengths represent functionally weakly connected neurons (Figure 2C). Finally, we defined each cortical core ensemble as a group of neurons that can be used to predict each visual stimulus with higher performance (AUC) and have high node strength (Figure 2D).

**Figure 2.**
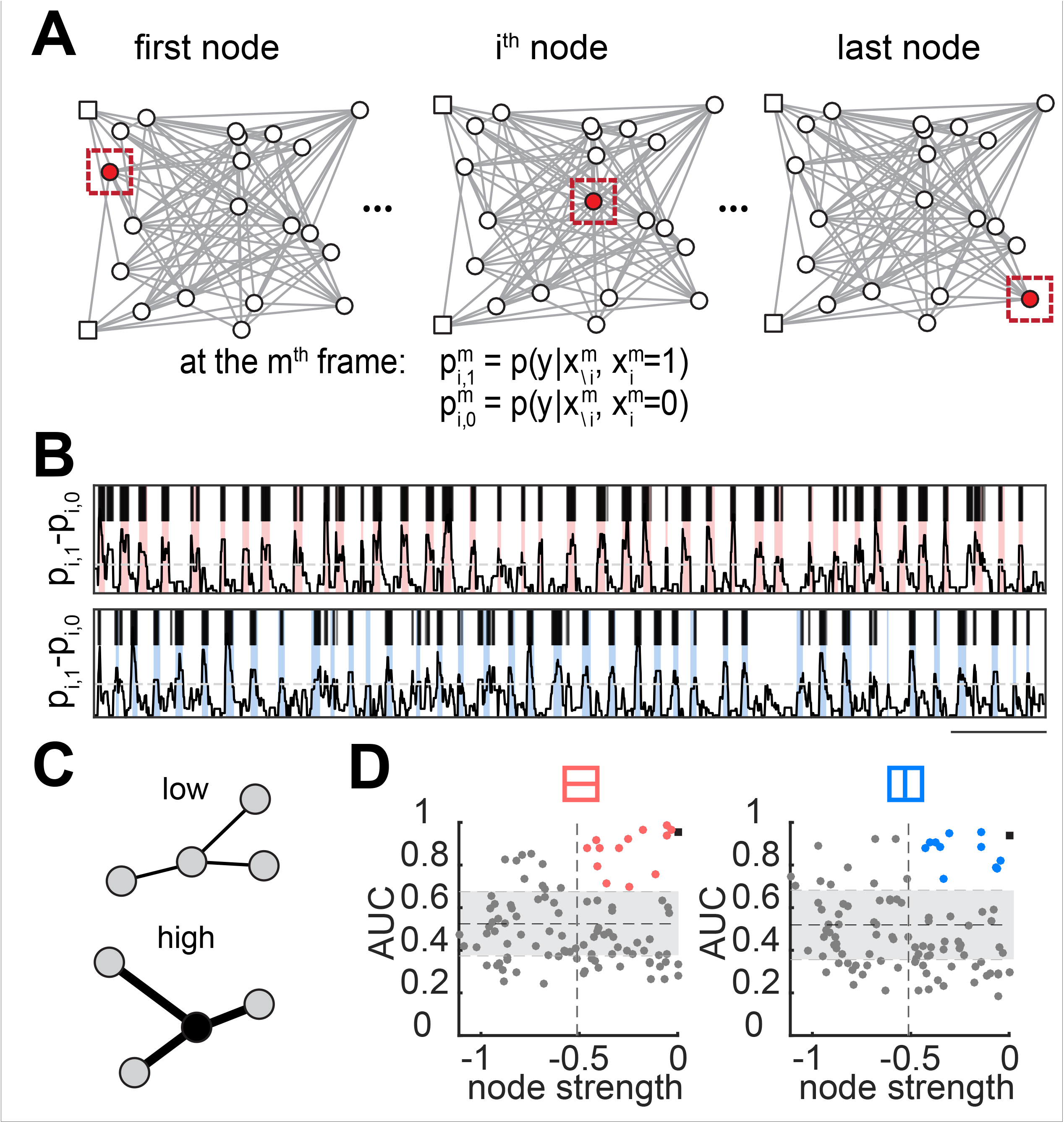
Identification of core neurons in cortical ensembles from CRFs. **A.** Schematic representation of core ensemble extraction from CRF models with added nodes for different visual stimuli. The activity of the *i*^th^ neuron is set to ‘1’ or ‘0’ at each frame, and the log likelihood 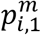 and 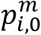 of modified population vectors is calculated. **B.** Log likelihood inference and prediction for cortical core ensembles representing horizontal (top; red) or vertical (bottom; blue) drifting-gratings. **C.** Graphical representation of node strength magnitude. **D.** Core neurons from cortical ensembles for two visual stimuli defined as neurons with high AUC and high node strength (top right quadrant). Confidence levels were defined from CRF models of shuffled data (grey bars). Related to Figure S1.

To demonstrate the neurophysiological meaning of cortical core ensembles we analyzed publically open datasets (Allen Brain Observatory Release 2016) that contain data from layer 2/3 of primary visual cortex consisting in several visual stimuli types with different experimental settings. Cortical core ensembles extracted from CRF models were able to predict several different orientations from visual stimuli in layer 2/3 of primary visual cortex (Figure 3A). Even more, cortical core ensembles revealed that classification performance to different drifting-gratings was significantly better for lower temporal frequencies, as measured by ROC curves and AUC values (Figures 3B-D; classification AUC: AUC_TF_=1 = 0.9129 ± 0.0134, AUC_TF_=2 = 0.8973 ± 0.0141, AUC_TF_=4 = 0.8063 ± 0.0226, AUC_TF_=8 = 0.7045 ± 0.0262, AUC_TF_=15 = 0.6517 ± 0.0171; p_1,4_<0.01, p_1,8_, p_1,15_<0.001; n = 5 animals, 20 ensembles; Wilcoxon rank sum test).

**Figure 3.**
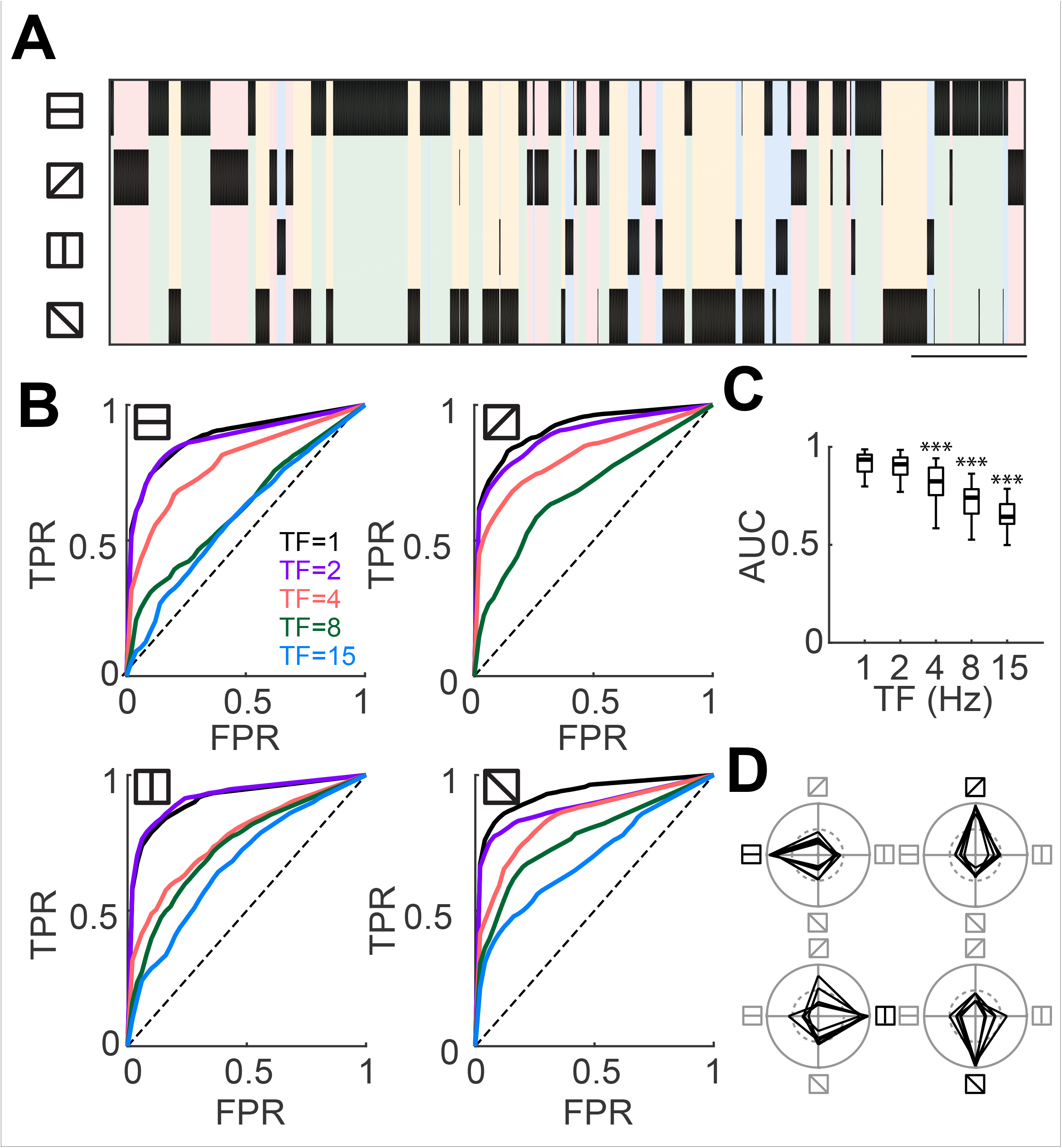
Cortical core ensembles have better predictive performance at low temporal frequencies. **A.** Temporal course of core ensembles extracted from CRFs. Colored stripes indicate different visual stimuli. Scale bar: 200 frames. **B.** ROC curves of core ensembles predicting different temporal frequencies (TF: 1, 2, 4, 8 and 15 Hz). Dashed line represents random classification performance. **C.** AUC for different TFs (classification AUC: TF 1 = 0.9129 ± 0.0134, TF 2 = 0.8973 ± 0.0141, TF 4 = 0.8063 ± 0.0226, TF 8 = 0.7045 ± 0.0262, TF 15 = 0.6517 ± 0.0171; p1,2=0.4249 n.s; p1,4=0.0003***; p1,8=2.9598e-07***; p1,15=6.7956e-08***). Note that cortical ensembles have better prediction performance for low TFs. **D.** Preferred orientation selectivity of core cortical ensembles for TF=1Hz. The radius of each circle depicts AUC values from zero (center) to 1 (border). Dotted inner circles represent random performance (AUC=0.5). Data presented as box and whisker plots displaying median and interquartile ranges (n = 5 mice, 20 ensembles; Wilcoxon rank sum test).

### Efficiency of cortical core ensembles

We next investigated whether core ensembles were optimal for predicting visual stimuli. To do so, we randomly shuffled population vectors containing the core neurons by adding or removing elements from the group, and examined the stimulus prediction performance. The similarity function and prediction performance had a maximum value when the size of core ensembles was unchanged (Figures 4A-C; similarity: 0.2887 ± 0.0926 [S.D.]; prediction: AUC 0.9383 ± 0.0333). We also calculated three standard measurements from the number of true positives (TP), true negatives (TN), false positives (FP) and false negatives (FN): accuracy, defined as (TP+TN)/(TP+TN+FP+FN); precision, defined as TP/(TP+FP); and recall, defined as TP/(TP+FN). Using these metrics, we demonstrated that core ensembles achieve the best accuracy, precision and recall when predicting visual stimuli, compared with resized ensembles (Figures S2A-C; accuracy: 0.8367 ± 0.0623, precision: 0.6175 ± 0.1752, recall: 0.9067 ± 0.0717).

**Figure 4.**
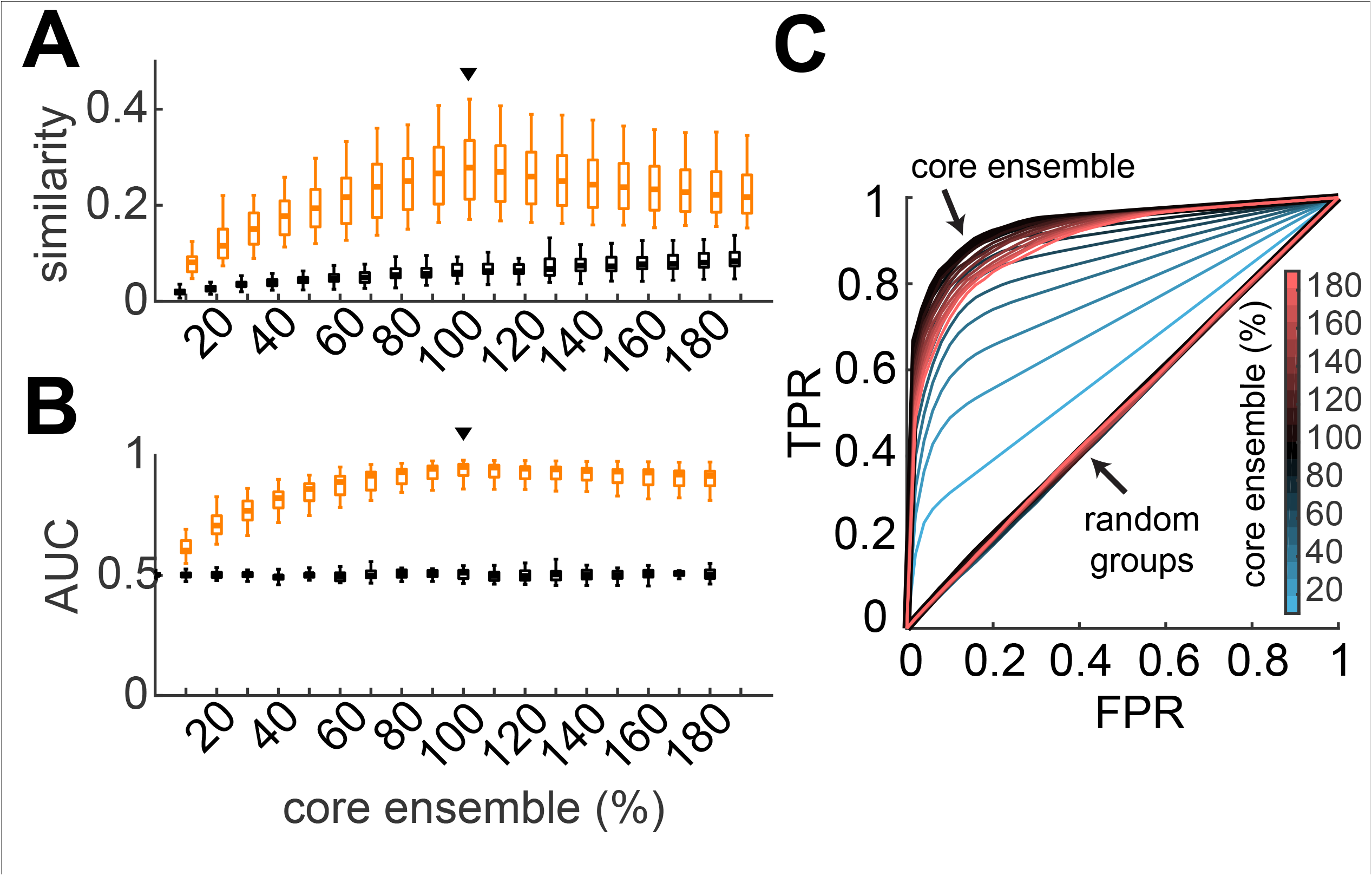
Efficacy of cortical core ensembles. **A**. Cosine similarity and **B**. AUC values from core ensembles extracted from CRFs that have been randomly down-sampled or up-sampled (orange). Randomly chosen neuronal groups are depicted in black. Black triangle indicates the original core ensemble size. Note that randomly removing or adding elements from the core ensemble decreases the ability to predict visual stimuli. **C**. ROC curves of randomly down-sampled or upsampled core ensembles, and randomly chosen neuronal groups. Line color represents the size of ensembles as a ratio with the original core ensemble size. (n=6 mice, 20 ensembles; Wilcoxon rank sum test). Related to Figure S2.

These results showed that cortical core ensembles extracted from CRFs represent an efficient population to predict external visual stimuli. This raises the question of whether such core ensembles are a specific non-random subgroup. To answer this, we randomly sampled a subset from all the population of neurons ranging from 10% to 190% of the total size of core ensembles. We observed that the prediction performance from random groups of neurons was significantly lower than the core ensembles extracted from CRF models (Figures 4A-C, black; Figures S2A-C, black; similarity: 0.1985 ± 0.0590 [S.D.], AUC: 0.5029 ± 0.0616, accuracy: 0.6906 ± 0.0671, precision: 0.2570 ± 0.1756, recall: 0.3617 ± 0.1027), indicating that cortical core ensembles achieved higher classification performance than random sets of neurons.

### Identification of neurons with pattern completion capability using CRFs models

One major advantage of CRFs is their ability to construct graphical models that capture network properties that could be used to target individual neurons. We therefore decided to exploit this advantage by targeting neurons with optogenetics in order to alter circuit function. It has been recently shown that the repetitive activation of an identified neuronal population with two-photon optogenetics imprints an artificial cortical ensemble that can be recalled later on by activating specific members of the ensemble (Carrillo-Reid et al., 2016). Since CRFs can be used to identify core neurons from cortical ensembles evoked by visual stimuli, we hypothesize that in the case of artificially imprinted cortical ensembles core neurons extracted from CRFs would represent neurons with pattern completion capability. In order to investigate that, we used simultaneous two-photon imaging and two-photon optogenetic experiments with single cell resolution (Figure S3A) to identify neurons with high node strength and high AUC values (Figure S3B). Our approach demonstrated as proof-of-principle that single-cell two-photon optogenetic stimulation of identified neurons (AUC: 0.8397 ± 0.0361; node strength: - 0.1405 ± 0.0770) was able to evoke pattern completion of artificially imprinted ensembles, whereas single cell stimulation of neurons with low node strength or low AUC values (AUC: 0.5680 ± 0.0292; node strength: -1.0332 ± 0.0573) was unable to recall artificially imprinted cortical ensembles (Figures S3C-D). These experiments support the hypothesis that neurons with pattern completion capability have high functional connectivity with other members of artificially imprinted cortical ensembles (Carrillo-Reid et al., 2016).

### CRFs models reveal the reconfiguration of cortical microcircuits

Another advantage of CRF models could be the ability to capture network changes in cortical ensembles under different experimental conditions. To test this we compared CRF models estimated from data before and after two-photon population manipulation of a given set of neurons for several times (Figure 5A), an experimental protocol that reconfigures network activity building new coactive ensembles (Carrillo-Reid et al., 2016). To visualize network changes in neurons with pattern completion capability we constructed isomorphic graphs from the CRF models and arranged them in a circular visualization before and after optogenetic manipulation (Figure 5B). After the artificial ensemble was imprinted, we observed that neurons with pattern completion capability increased their predictive performance and node strength (Figure 5C; stimulated neuron: node strength increased from: -0.1539 to 0.1609, AUC increased from: 0.5234 to 0.7447). This demonstrated as proof-of-principle that CRF models can be used to study network of specific neurons and those very CRFs can also be used to target single neurons that play a key role in the computational properties of cortical microcircuits. Interestingly, for non-photostimulated neurons, the graph properties of CRFs before and after population photostimulation remained stable (Figure S4) suggesting that imprinted ensembles have been added to cortical microcircuits while preserving a balance with the overall network structure.

**Figure 5.**
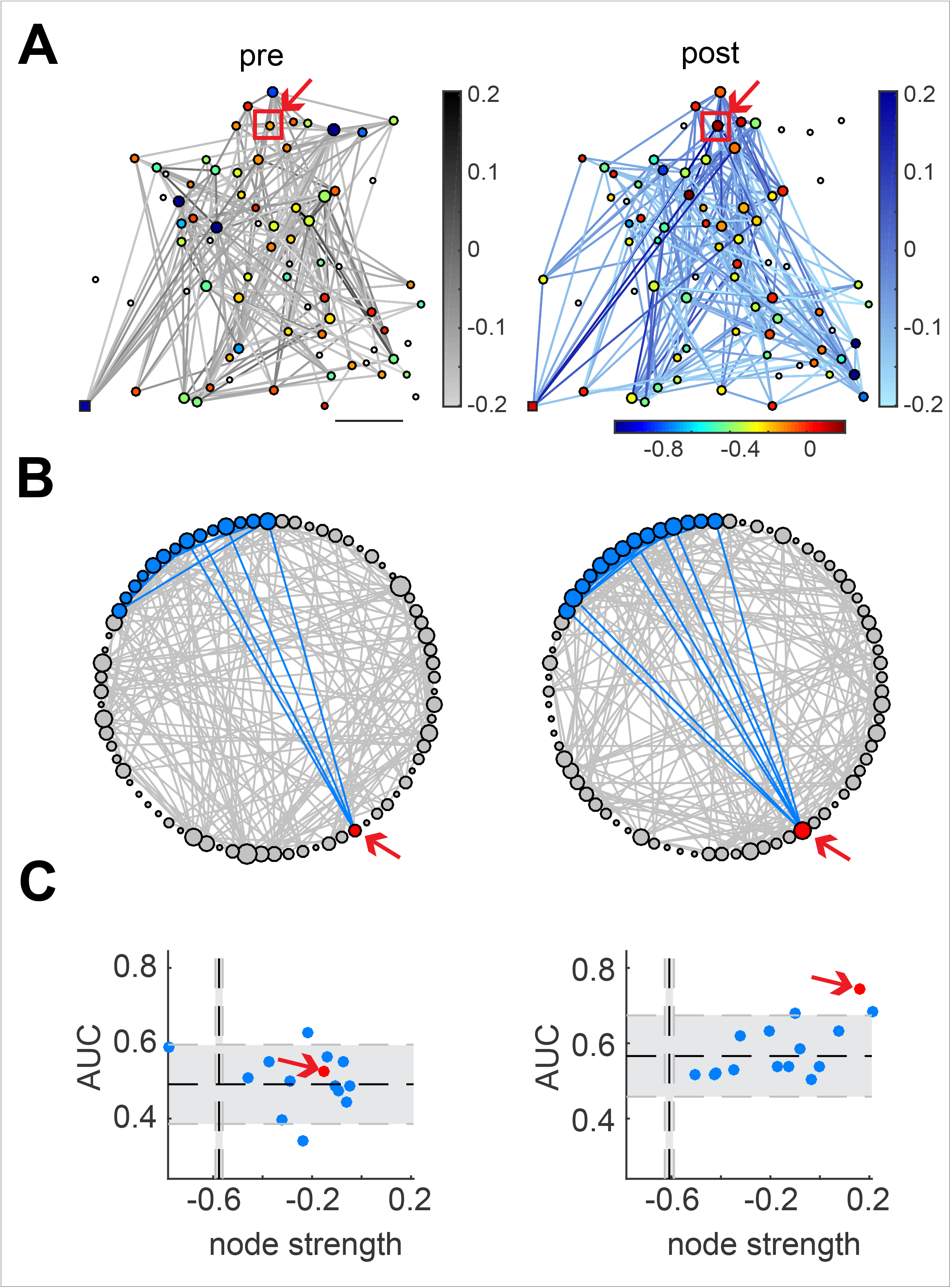
Reconfiguration of cortical microcircuits by two-photon optogenetic stimulation. **A**. Graphical models obtained using CRFs from simultaneous two-photon imaging and two-photon optogenetic stimulation of a neuron with pattern completion capability (Related to Figure S3) before (left) and after (right) two-photon optogenetic ensemble imprinting. Square on bottom left represents added node for optogenetic stimuli. Edge color tone represents edge potential strength (*φ*_11_); node color represents node strength. Node size represents node degree. Scale bar: 50μm. **B**. Isomorphic graphs of CRFs models before (pre) and after (post) ensemble imprinting. Blue neurons were stimulated with two-photon optogenetics (60 trials; 4Hz). Connections between photostimulated neurons are shown in blue. Red dot represents stimulated neuron (arrow). **C.** Node strength and AUC values showed network changes of neurons with pattern completion capability. The stimulated neuron is represented in red before (left) and after (right) ensemble imprinting. Confidence levels calculated from random data are depicted by grey bars. Related to Figures S3 and S4.

## Discussion

### Machine learning analysis of functional connectivity in cortical microcircuits

In this study, we provide a novel tool for the identification and targeting of key members of cortical ensembles from population calcium imaging recordings of mouse primary visual cortex *in vivo* using Conditional Random Fields (CRFs), a probabilistic and graphical machine learning method. As opposed to traditional descriptive approaches for neuronal ensemble identification and network analyses based on correlations between pairs of neurons, machine learning methods represent an empirically grounded approach to create models that aim to capture the functional structure of neuronal circuits and also provide information about the network properties of individual neurons within a population.

In the past decades, graph theory has been applied to characterize the structure and function of neuronal networks (Achard and Bullmore, 2007; Bettencourt et al., 2007; Chiang et al., 2016; Downes et al., 2012; Fair et al., 2008; Hagmann et al., 2008; Iturria-Medina et al., 2008; Oh et al., 2014; Supekar et al., 2008; Yu et al., 2008; Zuo et al., 2012). While most of these studies operated on functional recordings across multiple brain regions (Achard and Bullmore, 2007; Chiang et al., 2016; Fair et al., 2008; Hinne et al., 2013; Zuo et al., 2012), only a few have focused on the general network properties of cortical circuits with recordings from single neurons (Bonifazi et al., 2009; Sadovsky and MacLean, 2014; Stetter et al., 2012; Yatsenko et al., 2015). The majority of methods applied to infer network properties in brain slices (Cossart et al., 2003; Ikegaya et al., 2004; Mao et al., 2001; Sadovsky and MacLean, 2014; Stetter et al., 2012) or *in vivo* (Yatsenko et al., 2015) operate on the correlation matrix, and aim to recover the functional dependencies between observed neurons. However, these methods are model-free, therefore are incapable of describing the overall network dynamics based on the probability distribution of neuronal ensembles. Our method provides an alternative by directly modeling the statistical dependencies of each neuron. The generation of a graphical model of the circuit has obvious advantages, since it provides a direct link to its functional connectivity structure, enabling the design of close-loop experiments to target core neurons during behavioral tasks.

### CRFs graphical models identify neuronal ensembles

Compared with fully generative models such as Markov Random Fields and Bayesian Networks that make assumptions on the dependencies between all the observed variables from the model, CRFs only model the hidden system states dependent on observed features. Since no independence assumptions are made between observed variables, CRFs avoid potential errors introduced by unobserved common inputs. Additionally, given the finite number of network states described by population activity, the conditional distribution is sufficient for making predictions, both for the population state and for identifying core neurons in each state. Compared with other discriminative finite-state models such as Maximum Entropy Markov Models (MEMM), CRFs use global normalizers to overcome the local bias in MEMM induced by local normalizers, and have been shown to achieve higher accuracy in diverse applications (Lafferty et al., 2001). Therefore, CRFs appear to be promising for modeling cortical functional connectivity and for identifying core ensembles that could be easily manipulated by two-photon optogenetics.

The computational difficulty in constructing CRFs lies in recovering the global normalizer (the partition function) and gradients of global normalizer. With an arbitrary graph structure, this problem is often intractable. But recent advances that combines Bethe free energy approximation and Frank-Wolfe methods for inference and learning model parameters allow fast and relatively accurate construction of cyclic CRFs (Tang et al., 2016). Thus, CRFs can in principle be applied to datasets with thousands of interconnected neurons. However, for datasets with more neurons (and therefore more random variables and larger networks), CRFs (like most machine learning approaches) would require an increasingly large number of samples in the training dataset.

### Advantages and limitations

The key advantage of CRF models is their ability to model the circuit explicitly, in a manner than can be used for targeted manipulation. In visual cortex, non-targeted electrical stimulation has been used for decades as an attempt to provide useful visual sensations to patients that have lost the functionality of their eyes (Brindley and Lewin, 1986). The sensations produced by electrical stimulation of the visual cortex were termed phosphenes since they represented bright spots. To improve prostheses, one could in principle train patients using devices with a large number of electrodes (Shepherd et al., 2013). Our results suggest that after a given network have been imprinted (Carrillo-Reid et al., 2016), the identification of neurons with pattern completion capability could be used to reduce the number of active points that require stimulation and pinpoint them with surgical accuracy. The further development of network models based on population activity that can predict a given set of features embedded in visual stimuli will be crucial for the efficient manipulation of cortical ensembles.

We previously showed that population vectors defining a group (i.e. a cortical ensemble) can be extracted from multidimensional arrays by performing singular value decomposition (SVD) (Carrillo-Reid et al., 2015a; Carrillo-Reid et al., 2015b). Even though SVD can identify cortical ensembles reliably, it lacks a structured graphical model that allows the systematic study of changes in network properties.

One limitation for the current CRF learning algorithms is the computation of a sparse graph structure which is less prone to over-fitting of the data. Certainly, due to some modeling assumptions, computational considerations and the finiteness of training data, the learned graphical structure and parameters may not be the globally best probabilistic model of the data. However, it is recovered efficiently rather than requiring an exhaustive and unrealistic exploration of all possible structures and parameter combinations. Additionally, approximations during the parameter learning step can sometimes compromise the global optimality guarantees.

Finally, it has been shown that the connectivity of diverse systems described by graphs with complex topologies follow a scale-free power-law distribution (Barabasi and Albert, 1999). Scale-free networks are characterized by the existence of a small subset of nodes with high connectivity (Carrillo-Reid et al., 2015a). Similarly, cortical core ensembles described by CRFs could be characterized by a subset of neurons with strong synaptic connections. The existence of neurons with pattern completion capability has been demonstrated in previous studies where perturbing the activity of single neurons was able to change the overall network dynamics (Bonifazi et al., 2009; Carrillo-Reid et al., 2016; Hagmann et al., 2008). However, the efficient identification of such neurons from network models that capture the overall properties of neuronal microcircuits was lacking.

Our results suggest that the structural and predictive parameters defined by CRFs models could be used in the design of closed-loop experiments with single cell resolution to investigate the role of a specific subpopulation of neurons in a given cortical microcircuit during different behavioral events.

## Experimental Procedures

### Animals and surgery

All experimental procedures were carried out in accordance with the US National Institutes of Health and Columbia University Institutional Animal Care and Use Committee and have been described previously (Carrillo-Reid et al., 2016). Briefly, simultaneous two-photon imaging and two-photon optogenetic experiments were performed on C57BL/6 male mice. Virus AAV1-syn-GCaMP6s-WPRE-SV40 and AVVdj-CaMKIIa-C1V1(E162T)-TS-P2A-mCherry-WPRE were injected simultaneously into layer 2/3 of left primary visual cortex (2.5 mm lateral and 0.3 mm anterior from the lambda, 200 μm from pia). After 3 weeks mice were anesthetized with isoflurane (1-2%) and a titanium head plate was attached to the skull using dental cement. Dexamethasone sodium phosphate (2 mg/kg) and enrofloxacin (4.47 mg/kg) were administered subcutaneously. Carprofen (5 mg/kg) was administered intraperitoneally. After surgery animals received carprofen injections for 2 days as post-operative pain medication. A reinforced thinned skull window for chronic imaging (2 mm in diameter) was made above the injection site using a dental drill. A 3-mm circular glass coverslip was placed and sealed using a cyanoacrylate adhesive (Drew et al., 2010). Imaging experiments were performed 7~28 days after head plate fixation. During recording sessions mouse is awake (head fixed) and can move freely on a circular treadmill.

### Visual Stimulation

Visual stimuli were generated using MATLAB Psychophysics Toolbox and displayed on a LCD monitor positioned 15 cm from the right eye at 45° to the long axis of the animal. Population activity corresponding to two-photon stimulation of targeted neurons in layer 2/3 of visual cortex was recorded with the monitor displaying a gray screen with mean luminescence similar to drifting-gratings. The imaging setup and the objective were completely enclosed with blackout fabric and a black electrical tape. Visual stimuli consisted of full-field sine wave drifting-gratings (100% contrast, 0.035 cycles/°, 2 cycles/sec) drifting in two orthogonal directions presented for 4 sec, followed by 6 sec of mean luminescence.

### Simultaneous two-photon calcium imaging and photostimulation

Two-photon imaging and optogenetic photostimulation were performed with two different femtosecond-pulsed lasers attached to a commercial microscope. An imaging laser (λ = 940 nm) was used to excite a genetically encoded calcium indicator (GCaMP6s) while a photostimulation laser (λ = 1064 nm) was used to excite a red shifted opsin (C1V1) that preferentially responds to longer wavelengths (Packer et al., 2012).

The two laser beams on the sample are individually controlled by two independent sets of galvanometric scanning mirrors. The imaged field of view was ~240X240 μm (25X NA 1.05 XLPlan N objective), comprising 60-100 neurons.

We adjusted the power and duration of photostimulation such that the amplitude of calcium transients evoked by C1V1 activation mimic the amplitude of calcium transients evoked by visual stimulation with drifting-gratings. Single cell photostimulation was performed with a spiral pattern delivered from the center of the cell to the boundaries of the soma at 0.001 pix/ s for one second (Carrillo-Reid et al., 2016).

### Image processing

Image processing was performed with Image J (v.1.42q, National Institutes of Health) and custom made programs written in MATLAB as previously described. Acquired images were processed to correct motion artifacts using TurboReg . Neuronal contours were automatically identified using independent component analysis and image segmentation (Mukamel et al., 2009). Calcium transients were computed as changes in fluorescence: (F_i_ – F_o_)/F_o_, where F_i_ denotes the fluorescence intensity at any frame and Fo denotes the basal fluorescence of each neuron. Spikes were inferred from the gradient (first time derivative) of calcium signals with a threshold of 3 S.D. above noise level. We constructed an *N* x *F* binary matrix, where *N* denotes the number of active neurons and *F* represents the total number of frames for each movie. Peaks of synchronous activity describe population vectors (Carrillo-Reid et al., 2008).

### Population vectors

We constructed multidimensional population vectors that represent the simultaneous activation of different neurons., only high-activity frames were used. An activity threshold was determined by generating 1000 shuffled raster plots and comparing the distribution of the random peaks against the peaks of synchrony observed in the real data (Shmiel et al., 2006). We tested the significance of population vectors against the null hypothesis that the synchronous firing of neuronal pools is given by a random process (Carrillo-Reid et al., 2015a; Carrillo-Reid et al., 2015b; Shmiel et al., 2006). Such population vectors can be used to describe the network activity as a function of time (Brown et al., 2005; Carrillo-Reid et al., 2008; Sasaki et al., 2007; Schreiber et al., 2003; Stopfer et al., 2003). The number of dimensions for each experiment is given by the total number of active cells. The similarity index between a pair of vectors is then defined by their normalized inner product (Carrillo-Reid et al., 2008; Sasaki et al., 2007; Schreiber et al., 2003), which represents the cosine of the angle between two vectors. Neuronal ensembles are defined by the concomitant firing of neuronal groups at different times.

### Allen Institute Brain Observatory dataset

To demonstrate the general applicability of our approach we analyzed a publicly available dataset from the Allen Brain Observatory (http://observatory.brain-map.org/visualcoding) along with the SDK for extracting fluorescence and dF/F (http://alleninstitute.github.io/AllenSDK/) by Allen Institute of Brain Science. Spikes were detected by first low-pass filtering the dF/F traces, then a threshold of 5 S.D. above noise level on the first derivative of filtered dF/F. The experiments IDs used are: 511507650, 511509529, 511510650, 511510670, and 511510855.

### Conditional Random Fields

We constructed conditional random fields (CRFs) as previously published (Tang et al., 2016), using indicator feature vectors ***x*** = [*x*^1^,*x*^2^, …,*x^M^*], where *x^m^* ∈ *χ*, for each edge and node, and target binary population activity vectors ***y*** = [*y*^1^,*y*^2^, …,*y^M^*], where 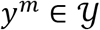, for *M* samples (time points). For each sample, the conditional probability can be expressed as:

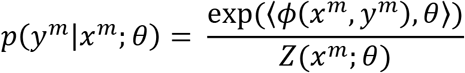

where *φ* is a vector of sufficient statistics of the distribution expressed in log-linear form, *θ* is a vector of parameters, and *Z* is the partition function:

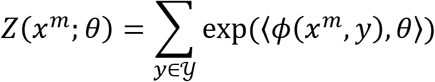

The conditional probability can be factored over a graph structure 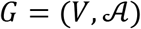, where *V* is the collection of nodes representing observation variables and target variables, and 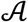 is the collection of subsets of *V*. The conditional dependencies can be then written as

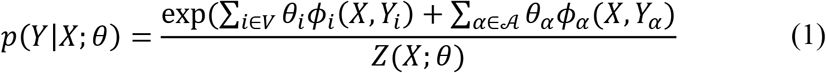

This model is a generalized version of Ising models, which have been previously applied to model neuronal networks (Yu et al., 2008). The log-likelihood of each observation can be then written as:

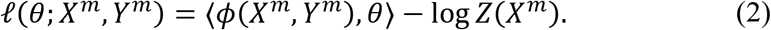

Given the inferred binary spikes from raw imaging data, we construct a CRF model by two steps: (1) structure learning, and (2) parameter learning. For structure learning, we learned a graph structure using *ℓ*_1_-regularized neighborhood-based logistic regression (Ravikumar et al., 2010):

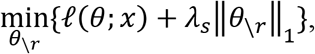

where

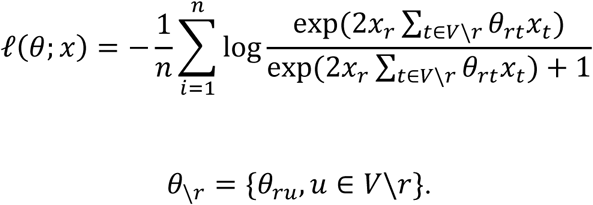

Here *λ_s_* is a regularization parameter that controls the sparsity (or conversely, the density) of the constructed graph structure. The final graph structure is obtained by thresholding the edge potentials with a given density preference *d*. Edges with potential values within the top *d* quantile are kept as the final structure. It is worth noting that although *d* could bias the result, varying *d* does not lead to density values that differ much. This is because of the sparsity induced by the *ℓ*_1_ regularizer.

Based on the learned structure, we use the Bethe approximation to approximate the partition function, and iterative Frank-Wolfe methods to perform parameter estimation by maximizing the log-likelihood of the observations (equation 2) with a quadratic regularizer (Tang et al., 2016):

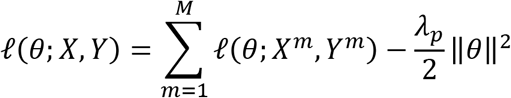

Here *λ_p_* is a regularization that controls the learnt parameters and helps preventing overfitting. Cross-validation was done to find the best *λ_s_*, *d* and *λ_p_* via held out model likelihood. We varied *λ_s_* with 6 values between 0.002 and 0.5, d with 6 values between 0.25 and 0.3, and *λ_p_* with 5 values between 10 and 10000, all sampled uniformly. To obtain the best model parameters, 90% data were used for training, while 10% data were withheld for cross-validation. The best model parameters were determined by calculating the likelihood of the withheld data and selecting the parameter set with a locally maximum likelihood in the parameter space.

### Node strength

We define the node strength as the sum of the ‘11’ term of edge potentials from all connecting edges:

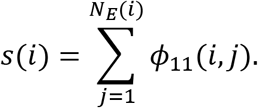

Here *N_E_(i)* denotes the number of connecting edges for node *i*. The defined node strength reflects the importance of a given cell in co-activating with other cells.

### Shuffling method

To generate shuffled models, we first randomize the spike raster matrices while preserving the activity per cell and per frame. Then, we trained CRF models using the shuffled spike matrices, with the cross-validated *λ_s_*, *d* and *λ_p_* from the real model. This procedure is repeated 100 times. Random level of node strength is determined by mean ± S.D. of mean node strengths from all shuffled models.

### Identifying the most representative cortical ensembles

To find the most representative cortical ensembles for each condition, we iterate through all the neurons and identify their contribution in predicting the stimulus conditions in the population. To this end, for the *i^th^* neuron in population, we set its activity to be ‘1’ and ‘0’ in turn, in all *M* frames. With the two resulting population vectors in the *m^th^* frame among all samples, we calculate the log-probability of them coming from the trained CRF model using equation (1):

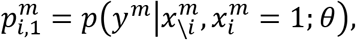

Then, we computed the log likelihood ratio:

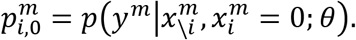

and calculated the standard receiver operating characteristic (ROC) curve with the ground truth as the timing of each presented visual stimuli. The prediction ability of all nodes for all presented stimuli is then represented by an area under curve (AUC) matrix *A*, where *A_i,s_* represents the AUC value of node *i* predicting stimulus *d*. Additionally, we calculated the node strength *S* = (*s_i_*) of each neuron in the CRF model.

We then computed prediction AUC *A^r^* and node strength *S^r^* of each node from 100 CRF models trained on shuffled data. The final core ensemble for stimulus *d* is defined as:

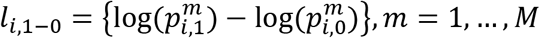

### Graph properties

Given the adjacency matrix *A* = (*a_υt_*) where *a_υ,t_* = 1 if node *υ* is linked to node *t*, we investigated the following graph properties: graph density, node degree, local clustering coefficient, and eigenvector centrality.

Graph density is calculated as the number of existing edges divided by the number of total possible edges:

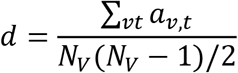

where *N_V_* is the number of vertices in the graph.

Node degree is defined for node *v* as the number of edges connected to it:

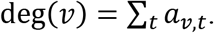

Local clustering coefficient is defined for each node as the fraction edges connected to it over the total number of possible edges between the node’s neighbors (nodes that have a direct connection with it).

Eigenvector centrality is defined on the relative centrality score matrix *X* = (*x_v_*), where

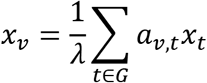

This can be written in the form of eigenvector equation:

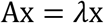

Solving the above equation gives a set of eigenvalues *λ* and associated eigenvectors. The *V^th^* entry of the eigenvector associated with the largest *λ* gives the eigenvector centrality for the *V^th^* node.

### Statistical Analysis

CRF models were trained using the Columbia Yeti shared HPC cluster. MATLAB R2016a (MathWorks) was used for data analysis. Statistical details of each specific experiment can be found in figure legends. All numbers in the text and figure legends denote mean ± S.E.M. unless otherwise indicated.

### Resource Availability

Code used in this paper can be found at https://github.com/hanshuting/graph_ensemble.

## Acknowledgments

We thank Kui Tang for CRFs code, laboratory members for valuable comments and virus injections, Columbia Yeti shared High Performance Computing Cluster for computation resources, Stanford Neuroscience Gene Vector and Virus Core for AAVdj virus and the NEI (DP1EY024503, R01EY011787) for funding. This material is based upon work supported by the Defense Advanced Research Projects Agency (DARPA) under Contract No. HR0011-17-C-0026 and SIMPLEX N66001-15-C-4032 and in part by the U. S. Army Research Laboratory and the U. S. Army Research Office under contract number W911NF-12-1-0594 (MURI). S.H. is a Howard Hughes Medical Institute International Student Research fellow. The authors declare no competing financial interests.

## Author Contributions

L.C.-R & S.H. contributed equally to this work. Conceptualization, L.C.-R, S.H, T.J. and R.Y.; Methodology, L.C.-R., S.H., E.T. & T.J.; Investigation, L.C.-R. & S.H.; Writing – Original Draft, L.C.-R. & S.H.; Writing – Review & Editing, L.C.-R., S.H., T.J., & R.Y.; Resources, L.C.-R., S.H. & T.J.; Funding Acquisition, T.J. & R.Y.

